# Towards a standardization of non-symbolic numerical experiments: GeNEsIS, a flexible and user-friendly tool to generate controlled stimuli

**DOI:** 10.1101/2021.03.03.433737

**Authors:** Mirko Zanon, Davide Potrich, Maria Bortot, Giorgio Vallortigara

## Abstract

Several studies have suggested that vertebrate and invertebrate species may possess a *number sense*, i.e. an ability to process in a non-symbolic and non-verbal way the numerousness of a set of items. However, this hypothesis has been challenged by the presence of other non-numerical continuous physical variables, that vary along with numerosity (e.g. any change in the number of visual physical elements in a set naturally involves a related change in visual features such as area, density, contour length and convex hull of the stimulus). It is therefore necessary to control and manipulate the continuous physical information when investigating the ability of humans and other animals to perceive numerousness. During decades of research, different methods have been implemented in order to address this issue, which has implications for experiments replicability and inter-species comparisons, since no general standardized procedure is currently being used. Here we present the “Generation of Numerical Elements Images Software” (GeNEsIS) for the creation of non-symbolic numerical arrays in a standardized and user-friendly environment. The main aim of this tool would be to provide researchers in the field of numerical cognition with a manageable and precise instrument to produce visual numerical arrays controlled for all the continuous variables; additionally, we implemented the possibility to actively guide stimuli presentation during habituation/dishabituation and dual-choice comparison tasks used in human and comparative research.

## Introduction

The ability to operate with numerical information is an essential skill to deal with our everyday life. However, the symbolic numerical system and the formal arithmetic developed by humans are believed to be rooted in a more ancient and possibly innate numerical ability, the so-called “*number sense”* (Dehaene, 1997). This sense of number is a non-symbolic, language-independent and evolutionary conserved ability that allows humans and other animals to represent numerical sets of physical items in the surrounding environment (Ferrigno & Cantlon, 2017; Vallortigara, 2014, 2017). Animals, both vertebrates and invertebrates, can take advantage of this ability in order to optimize foraging decision (vertebrates: Garland et al., 2012; Gazzola et al., 2018; Hanus & Call, 2007; Stancher et al., 2015; invertebrates: Bar-Shai et al., 2011; Hemptinne et al., 1992; Nelson & Jackson, 2012), conflicts between groups, defensive strategies (vertebrates: McComb et al., 1994; Potrich et al., 2015; Wilson et al., 2002; invertebrates: (Tanner, 2006), mating competition (vertebrates: Flay et al., 2009; invertebrates: Carazo et al., 2009) and parental care (Lyon, 2003).

It has been hypothesized that the *number sense* is actually instantiated in an “Approximate Number System” (ANS), a non-verbal mechanism that allows to estimate and represent numerosity of sets of physical elements, which obeys the Weber’s Law (Gallistel, 1990), i.e. as the ratio between the numbers to be discriminated increases, response times increases and accuracy decreased (e.g., discriminate 5 *vs*. 15, with a 0.33 ratio, is easier than discriminate 10 *vs*. 20, with a 0.5 ratio; Gallistel & Gelman, 1992; for general reviews see e.g. Butterworth, 1999; Hyde, 2011; Nieder & Dehaene, 2009). Evidence for a dedicated neural network for processing numerical information involving parietal and prefrontal cortical regions has been reported in humans (Arsalidou & Taylor, 2011; Piazza & Eger, 2016) and non-human primates (Wang et al., 2015). Furthermore, neurons that show tuned selectivity to specific numerousness have been described in both humans (see Nieder, 2016 for review) and non-human species, e.g. primates (Viswanathan & Nieder, 2013) and corvids (Ditz & Nieder, 2015; Wagener et al., 2018). Recently, evidence for neurons sensitive to numerousness has been reported in the dorso-central pallium of zebrafish (Messina et al., 2020a, 2020b).

The evidence supporting non-symbolic numerical estimation in a variety of species is robust; however, specifying the precise nature of what is actually estimated is challenging, since it has been argued that estimation may be guided by non-numerical continuous physical variables, such as the amount of stimulus extension in space (Leibovich et al., 2017). The debate regarding the existence of a *number sense* is rooted in a methodological issue for which it is empirically impossible to take apart numerical information from all other continuous properties at once, making it difficult to study the non-symbolic numerosity processed in isolation from continuous magnitudes. In a natural environment, a change in numerosity of a set of items usually involves a change in the overall items’ properties (such as volume, area and perimeter) that co-varies with numbers: five objects normally occupy a larger space than three objects. At the same time, if the elements are displaced at the same distance one from the others, the global occupied field (known also as convex-hull) will be different, and inversely, by pairing the global field, the larger group will present a higher density.

As an alternative to the *number sense theory*, some authors have suggested that numbers are estimated and compared by combining the different sensory cues comprising the numerical value, using a sensory-integration-system (Gebuis et al., 2016). According to another hypothesis numerosities and magnitudes would be processed holistically, thus arguing that it would be more appropriate to refer to a *sense of magnitude* than to a *sense of number* (Leibovich et al., 2017). Evidence supporting a broader processing of numerosity and continuous magnitudes have been reported by several studies. Using an algorithm capable to generate dot arrays in which continuous variables might be congruent or incongruent with the number of elements in the set, Gebuis and Reynvoet (2011, 2012) showed that participants’ judgement was influenced by the visual properties of the elements’ arrays. Continuous magnitude appears to influence performance even with numerosities in the subitizing range (Leibovich et al., 2015; Salti et al., 2017). Studies carried out in humans and different animal species have shown that numerical tasks might be influenced by non-numerical cues such as the overall surface area (Feigenson et al., 2002; Stevens et al., 2007; Tokita & Ishiguchi, 2010), the individual size (Beran et al., 2008; Henik et al., 2017), the spatial frequency (Felisatti et al., 2020), and the density of the elements array (Dakin et al., 2011; Gómez-Laplaza & Gerlai, 2013).

This debate between *sense of number* and *sense of magnitude* is leading researchers to pay particular attention to the control of continuous physical variables. The simultaneous balancing of all these variables at the same time, however, is not viable: for instance, when the convex hull of the stimuli increases, the density decreases and *vice versa*; similarly, when the overall area of two sets of elements with different numerousness is balanced, their overall contour length would differ (Gebuis et al., 2014; Leibovich & Henik, 2014; Mix et al., 2002). It is therefore important to set up control conditions in which the different continuous physical variables are randomized and contrasted with numerousness as such to rule out their possible use as non-numerical cues that may guide the discrimination.

The most common solution to the issue of continuous physical variables is to use sets of numerical stimuli controlled, in turn, for some (and different) visual features. The principle strategy is to make these continuous variables uninformative cues for discriminative judgement.

The use of computerized methods to create visual sets of stimuli allows reasonable advantages. In the last years, new algorithms have been developed for this purpose (see for example De Marco & Cutini, 2020; DeWind et al., 2015; Guillaume et al., 2020; Salti et al., 2017; see Table 1 for a comparison). These algorithms are usually based on strict mathematical constraints that tolerate a certain amount of error, usually minimal (e.g., up to ± 0.001%-pixel tolerance limit; De Marco & Cutini, 2020). Mathematical calculation also allows a standardized procedure to evaluate the continuous variables in the group of elements, thus the experimenter can study their impact on the subjects’ performance, as well as being provided with a general method that gives the possibility to study intra and inter-species differences. This minimizes the impact of biased errors and experimental procedure differences (De Marco & Cutini, 2020; Guillaume et al., 2020).

**Table 1:** Comparison between GeNEsIS and other programs developed until now. The green check symbol indicates the possibility for the program to perform the correspondent task, the red cross the absence of that tool. The yellow check indicates a limited possibility for the correspondent task: for instance, NASCO does not present the possibility to select a custom total perimeter value, but since in this software the radius is fixed inside an array, the perimeter could be indirectly controlled; in Salti’s software, instead, the controlled variables are not fully settable since the program allows to fix only their ratio across numerosities. With density we refer here to a measure of mean occupancy, as slightly differently defined in the correspondent papers (De Marco & Cutini, 2020; Gebuis & Reynvoet, 2012; Guillaume et al., 2020; Salti et al., 2017); while with inter distance we refer to the definition -most geometrically closer to distance-we gave (or similar definitions that can be found for example also in De Marco and Cutini (2020), even if only presented in output and thus not directly controlled).

|  | GeNEsIS | De Marco et al.<br>2020<br>(CUSTOM) | Guillaume et al.<br>2020 (NASCO) | Salti et al.<br>2017 | Gebuis et al.<br>2012 |
| --- | --- | --- | --- | --- | --- |
| Control over Convex Hull | ✓ | ✓ | ✓ | ✓ | ✓ |
| Control over Density | ✓ | ✓ | ✓ | ✗ | ✓ |
| Control over Inter Distance | ✓ | ✗ | ✗ | ✗ | ✗ |
| Control over Total Area | ✓ | ✓ | ✓ | ✓ | ✓ |
| Control over Total Perimeter | ✓ | ✓ | ✓ | ✓ | ✗ |
| Control over Radius | ✓ | ✓ | ✓ | ✓ | ✓ |
| Possibility to vary Radius between array's elements | ✓ | ✓ | ✗ | ✓ | ✓ |
| Possibility to fix minimum/maximum Radius | ✓ | ✗ | ✗ | ✗ | ✗ |
| Possibility to save images with a defined dimension in cm | ✓ | ✗ | ✗ | ✗ | ✗ |
| Possibility to create elements with a different shape | ✓ | ✗ | ✗ | ✗ | ✗ |
| Experiment presentation tool | ✓ | ✗ | ✓ | ✓ | ✓ |
| Graphic User Interface | ✓ | ✗ | ✓ | ✗ | ✗ |

An example of these recent tools is given by Guillaume et al. (2020); despite taking into careful account the influence of several continuous variables, such as area, size, convex hull, and element’s spatial occupancy, the program however does not allow a proper control on the internal size of elements, producing arrays composed of identical items. This may cause important differences in terms of spatial frequency.

An alternative tool was developed by De Marco and Cutini (2020), allowing a flexible creation of dot arrays; however, this program lacks some important features, such as an active control of the items’ inter distance, the minimum and maximum distance that can divide neighbour elements, as well as a fixed minimum and/or maximum dimension of elements. Besides, this program does not present an interactive interface that allows a non-expert user to simply interact with the program. The main goal of the algorithm we present here, GeNEsIS (*Generation of Numerical Elements Images Software*, GeNEsIS), would be to provide an alternative tool that integrates all the aforementioned features, helping in solving the issues related to the creation of numerical stimuli. The program will guide the user throughout the creation of sets of elements with a user-friendly interface, giving the possibility to freely utilize the output images for different purposes. Moreover, it provides a tool to perform the stimuli presentation on screen, throughout two of the most common experimental methods: a habituation/dishabituation task and a simultaneous dual-choice discrimination tasks. We believe that our software is a step towards a standardization of stimulus presentation in the study of numerical cognition. It improves the way in which experiments can be designed and controlled with respect to previous programs, adding more flexibility and a wider handling spectrum on continuous physical variables on which the experimenter could act (see Table 1 for a schematic comparison between the main existing programs, and Figure 5 for accuracy comparisons among the most recent ones).

**Figure 5.**
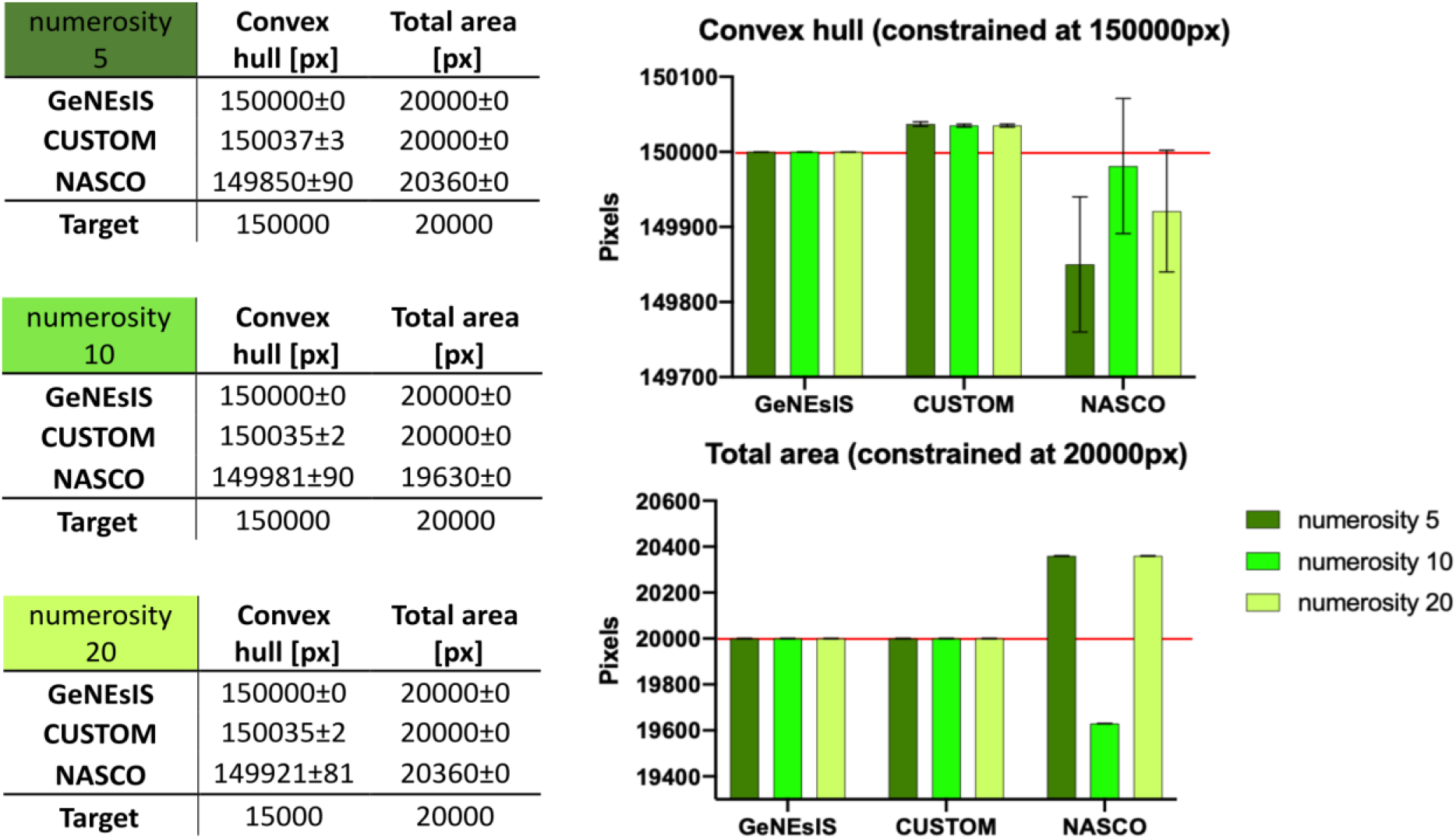
Comparison simulations between GeNEsIS and the most recent software (De Marco & Cutini, 2020; Guillaume et al., 2020). Three numerosities were generated (5,10 and 20), with 100 images per numerosity. The aim was to fix the convex hull (at 150000 px) and the total area (at 20000 px) across numerosities. The tolerance was set at very low levels to test the fine precision of the software (0.0001% or the minimum possible). Means with relative standard errors for the controlled variables are reported in the tables, and depicted in the bar graph as well, for a direct comparison.

### Materials and methods

We present a custom program (GeNEsIS) written in Matlab (Matlab R2019a, The MathWorks Inc., Natick, Massachusetts, USA), using *Appdesigner*. GeNEsIS is user-friendly, with a step-by-step tutorial; both the program code and the tutorial can be freely downloaded from https://github.com/MirkoZanon/GeNEsIS.

Using GeNEsIS, the user can create stimuli with completely custom characteristics, precisely controlling all the relevant parameters and physical variables like convex hull, density, mean inter-distance, total surface area and total perimeter (see Table 2 for an example of each controllable variable; program accuracy in a stimuli generation sample is shown in the Appendix I, see Supplementary Figure 1). GeNEsIS is easy to use and allows a large flexibility in the stimuli creation. Moreover, the program implements a tool to perform automatized experiments using habituation/dishabituation and simultaneous discrimination procedures; the script can also be adapted and modified to fulfil different user necessities in presenting stimuli in many different ways. The presentation on screen exploits *Psychtoolbox-3* (the Psychophysics Toolbox extensions; Brainard, 1997; Kleiner et al., 2007; Pelli, 1997). Noteworthy, the main tool to generate controlled stimuli is completely unrelated to any specific experiment or theory, thus it has a huge applicability spectrum and the creation of stimuli can be adapted to different theoretical frameworks.

**Table 2:** Summary of the main possible combinations between physical variables, as dictated by geometrical constraints. In particular, we consider different combinations of Inter Distance (ID), Density (D), Convex Hull (CH), Total Area (TA). Total Perimeter (TP), Radius (R) (this last can be free to vary or equal inside an array; R_min and R_max refers to the possibility to fix the minimum and maximum radius in the array). GeNEsIS can handle all these kinds of configurations, and even many other possible custom configurations that are less restrictive than these. The control of more variables at a time, different to the ones reported here, is not allowed given the geometrical limitations of these variables (i.e. their reciprocal dependency and their different covariation with numerosity).

|  | TA | TP |  |  |  |
| --- | --- | --- | --- | --- | --- |
| ID | TA+ID | TP+ID |  |  |  |
| D | TA+D | TP+D |  |  |  |
| CH | TA+CH | TP+CH |  |  |  |
|  | R_min | R_max | R_min+R_max | R_equal | R_free |
| TA+ID | TA+ID+R_min | TA+ID+R_max | TA+ID+R_min+R_max | TA+ID+R_equal | TA+ID+R_free |
| TA+D | TA+D+R_min | TA+D+R_max | TA+D+R_min+R_max | TA+D+R_equal | TA+D+R_free |
| TA+CH | TA+CH+R_min | TA+CH+R_max | TA+CH+R_min+R_max | TA+CH+R_equal | TA+CH+R_free |
| TP+ID | TP+ID+R_min | TP+ID+R_max | TP+ID+R_min+R_max | TP+ID+R_equal | TP+ID+R_free |
| TP+D | TP+D+R_min | TP+D+R_max | TP+D+R_min+R_max | TP+D+R_equal | TP+D+R_free |
| TP+CH | TP+CH+R_min | TP+CH+R_max | TP+CH+R_min+R_max | TP+CH+R_equal | TP+CH+R_free |

The program works with Matlab 2019 or more recent versions, and it has been largely tested, proving its functioning both in Windows PC (e.g Windows 10 Intel® Core™ i7-3770 - CPU: 3,40 GHz - RAM: 8 GB and Intel® Core™ i5-4570 - CPU: 3,2 GHz - RAM: 8 GB) and Mac (e.g MacBook Pro Intel® Core™ i7 - CPU: 2,6 GHz - SSD: 512 GB - RAM: 16 GB and Intel® Core™ i5 - CPU: 2,7 GHz - RAM: 8 GB).

### 1. The controlled physical variables

GeNEsIS allows complete control over six main continuous variables (i.e. inter distance, convex hull, density, radius, total perimeter and total area) and many other additional parameters (i.e., number of items in the array, shape and colour, arena type and dimension, accepted error and number of generations). Here we report a summary of all the parameters that the user can access and control, explaining how they are calculated.

- Inter-distance (ID): the average of the distances between all the possible pairs of elements (Figure 1a). These lasts are calculated as the euclidean distances between the element centres. ID is not defined for one element.
- Convex hull (CH): the area of the smallest convex polygon containing all the elements (Figure 1b); it is calculated through *convhull*() Matlab function using the points of all the elements’ perimeters. Thus, it can be seen as the surface area defined by the outermost stimuli perimeters. If only one element is presented, the CH value corresponds to its area.
- Density (D): the physical definition of elements’ density in a two-dimensional space, i.e. the number of elements (n) divided by the total occupied area (D = n/CH), (Figure 1c). D is not defined for one element.
- Radius: the specific dimension of the elements. For the circular shapes it is the proper circle radius (Figure 1d); for squares, diamonds and triangles it refers to the side dimension of the shapes.
- Total perimeter (TP): the sum of the perimeters (contours) of all elements (Figure 1e).
- Total area (TA): the sum of the areas (surfaces) of all elements (Figure 1f).

**Figure 1.**
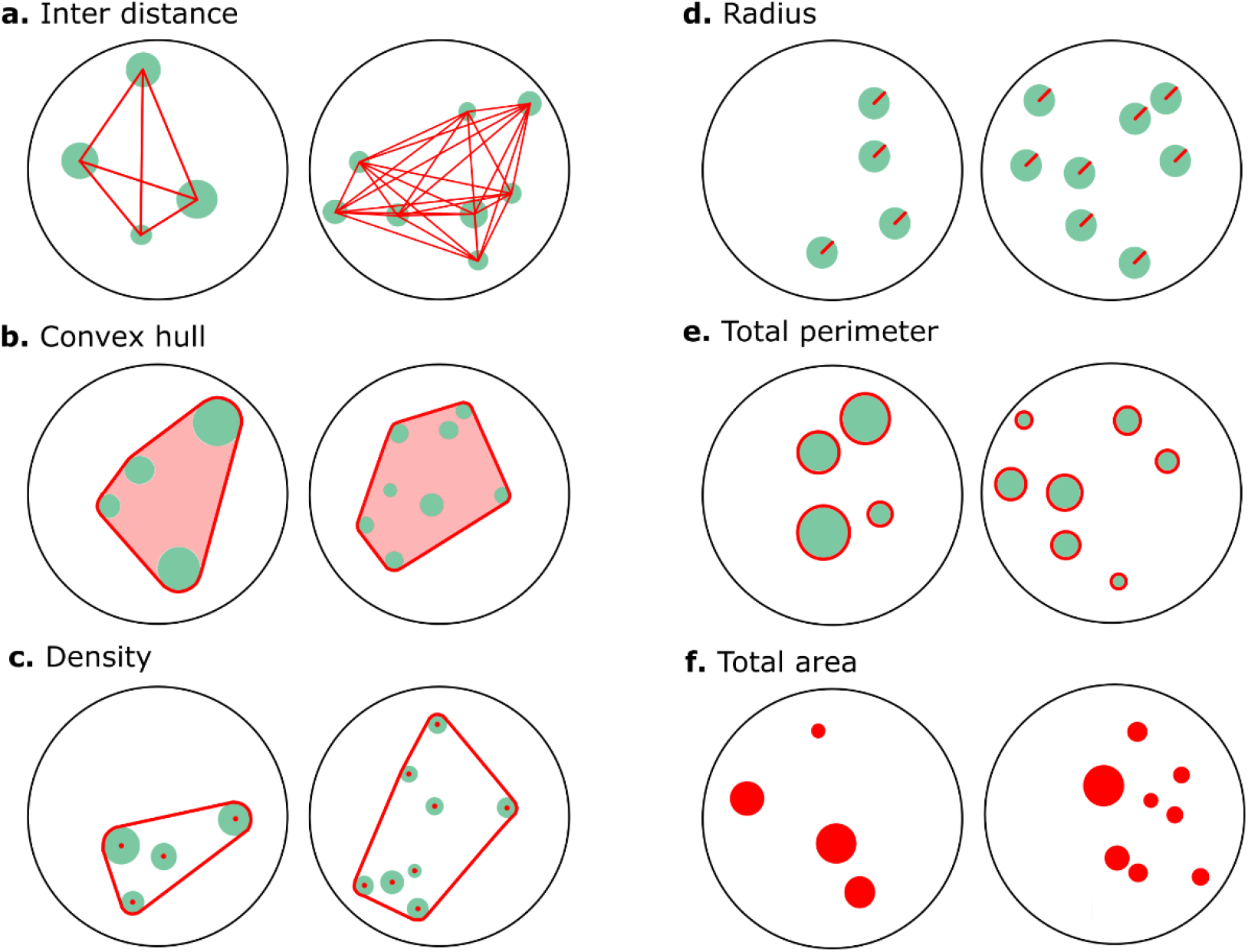
Example of stimuli with different numerosity (4 *vs*. 8 elements), controlled for the main continuous variables (sketched in red). **a)** Inter Distance (ID): both groups are constructed with the same mean inter distance between all elements; **b)** Convex Hull (CH): both groups are constructed with the same convex hull; **c)** Density (D): both groups are constructed with the same density (n/CH); **d)** Radius (R): both groups are constructed with all equal elements (fixed radius); **e)** Total Perimeter (TP): the sum of all the elements’ perimeters is the same for both groups; **f)** Total Area (TA): the sum of all the elements’ areas is the same for both groups. Combinations of one variable of the left group of controls (ID or CH or D) with one of the right (R or TP or TA) are also possible, allowing for the maximum flexibility (see also Table 2 and Figure 5).

Additional controls of secondary variables can be applied, concerning the following points.

- Number of elements: number of elements in the visual array (the user is allowed to select 1 to *n* possible elements).
- Arena radius: the dimension of the field in which elements are created; it is possible to choose a circular or squared arena (in which case the term ‘radius’ refers to half the square side). The colour can be set using the RGB colour code.
- Shape and colour: the characteristics of elements that can be created, choosing between circles, squares, rotated squares (diamonds), equilateral triangles. Completely custom colours can be set in RGB code.
- Accepted error: the tolerance of the program. Variables differing from the set value less than the ‘accepted error’ percent are considered as fulfilling the requirements.
- Generations: the number of stimuli that are created at a time.

### 2. GeNEsIS workflow

The program is structured in three steps: the computation of the stimuli with controlled physical variables (*GeNEsIS_create*), the saving of the final images with custom layout (*GeNEsIS_save*) and the optional presentation of images on screen for experimental use (*GeNEsIS_display*).

### First step: the computation of stimuli with controlled geometrical characteristics

This step is performed with *GeNEsIS_create*. Here the user will be guided through two passages (Figure 2): the control of elements’ distribution and of their shape. The first step allows the creation of stimuli distribution in space (controlling points’ convex hull, density, inter-distance), while the second step allows the control of area, perimeter, type of shape, and finally to eventually adjust the shaped elements’ distribution to match the selected convex hull (see later for a detailed discussion).

**Figure 2.**
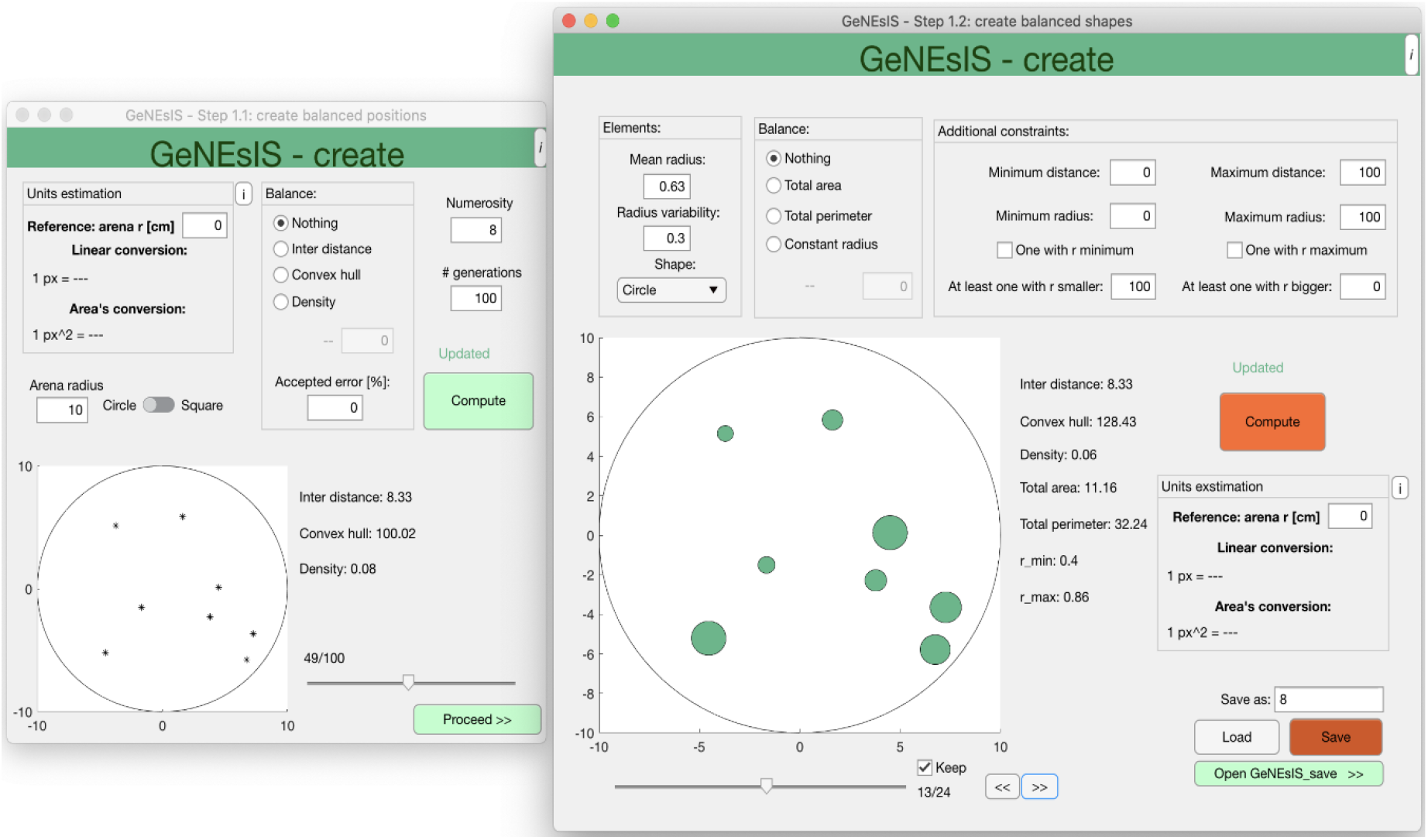
Graphic interface for the generation of GeNEsIS stimuli, (first step, GeNEsIS_create). In this step the user can choose the continuous variables to control (ID, CH, D in the first passage -left-; R, TP, TA, shape in the second passage -right-). The program has an intuitive user-friendly interface to guide through all the process of stimuli computation.

In the first passage, after setting the arena dimension and shape (i.e., circular or squared arena), and the number of elements, the script will continuously generate configurations with these characteristics, keeping only the ones fulfilling the spatial distribution constraints. It will stop after collecting the selected number of generations. Later, it will further discard the ones that do not meet the chosen shape/geometrical constraints (see below). For this reason, it would be desirable to perform a big number of generations in order to obtain a final pool of a reasonable number of items. Since this could take a long time, an *‘accepted error’* option is settable in order to keep also configurations differing from the set constraints by an error smaller than the accepted one. Already with an accepted error of 0.01%, there is a huge improvement in the computational time (for example, applying this threshold, in most of the cases there is an increase in speed of more than two times). As remarked, all the variables are here calculated for points’ distributions.

In the second passage, the geometrical characteristics of the elements like shape, dimensions, areas and perimeters can be set. It is also possible to impose strong constraints on the radii. If the elements have not a fixed radius, a mean one and a variability can be chosen. The program will try to generate the elements with radii belonging to a gaussian distribution with that mean and sigma (variability). Also, the minimum and maximum radius for the elements arrays are settable and can be maintained fixed across arrays (Figure 6). Every time the user changes the parameters and re-compute the elements, a new random set fulfilling the user’s constraints is generated on the same positions. A fundamental step to improve the program precision when controlling the CH is the refinement of the elements’ positions. If the user chooses a fixed CH, only the initial disposition of points (in the previous passage) will fulfil that value. However, when shaping the elements around these points, their extensions in space increase and so the effective CH. For this reason, the ‘real’ CH (i.e. considering the shapes’ perimeters) is recalculated in this second passage, and the elements’ positions are moved by small steps towards their centre of mass until the desired CH is matched. In this way GeNEsIS will give a more precise estimation of the CH (with a negligible error at the pixel level, related to the set tolerance), considering the physical extension of all the elements. If the user does not select a fixed CH, the program will simply calculate and update the effective CH value. Saving these data, a Matlab GeNEsIS file is created, ready to be used to customize layout characteristics, as reported in the second step.

**Figure 6.**
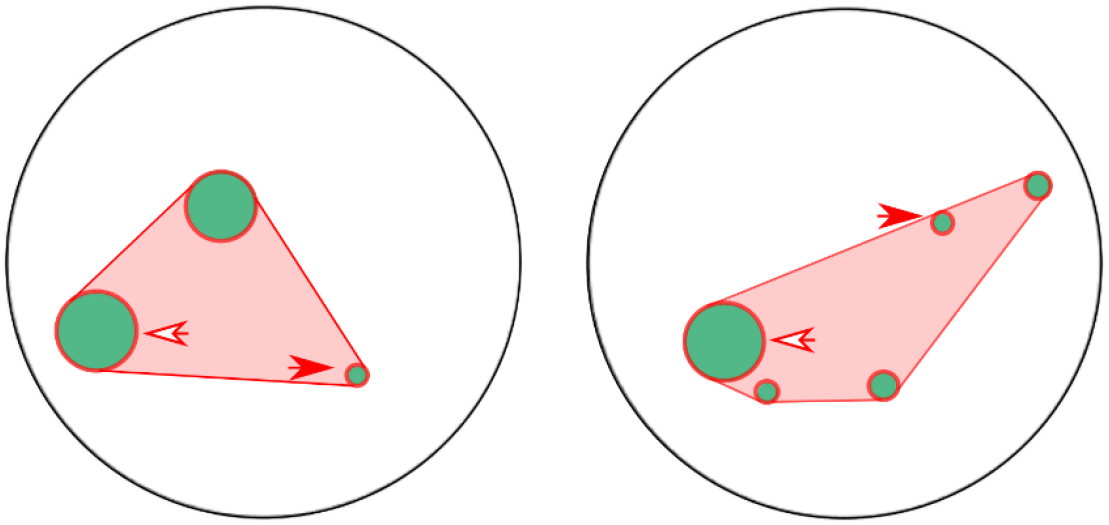
Example of stimuli with different numerosity 3 *vs*. 5 elements), controlled for a combination of different variables (sketched in red). In particular, both groups have the same convex hull and total perimeter. Moreover, a strict control over the radii is applied, imposing the same element size for the smallest and biggest items (i.e. the radii of the smallest elements are the same in both groups -red arrows-, that’s true for the biggest elements as well -white arrows).

### Second step: the saving of the final images with custom layout

This step is performed with *GeNEsIS_save*. After the creation of the stimuli pools, it is possible to save the created stimuli configurations as images ready to be presented during the experiment. Here, the effective dimensions, colours and other graphic characteristics can be chosen simultaneously for two different sets (Figure 3). The tool allows for the largest flexibility in colour customization, and moreover it gives the possibility to save different configurations of images, as single stimulus or couple of stimuli in different orientations (horizontal or vertical). The output can be a PNG image file or a Matlab matrix of images, ready to be used in an automated presentation with a computer. The elements’ dimensions in the output image can be precisely optimized, setting the output canvas dimension in pixels and the corresponding effective dimension in centimetres. In this way, fixing the desired effective dimension of the arena -taken as reference-, the user will obtain a figure with the exact final size. An excel file is also saved in output as reference, with all the relevant characteristics of all the generated stimuli.

**Figure 3.**
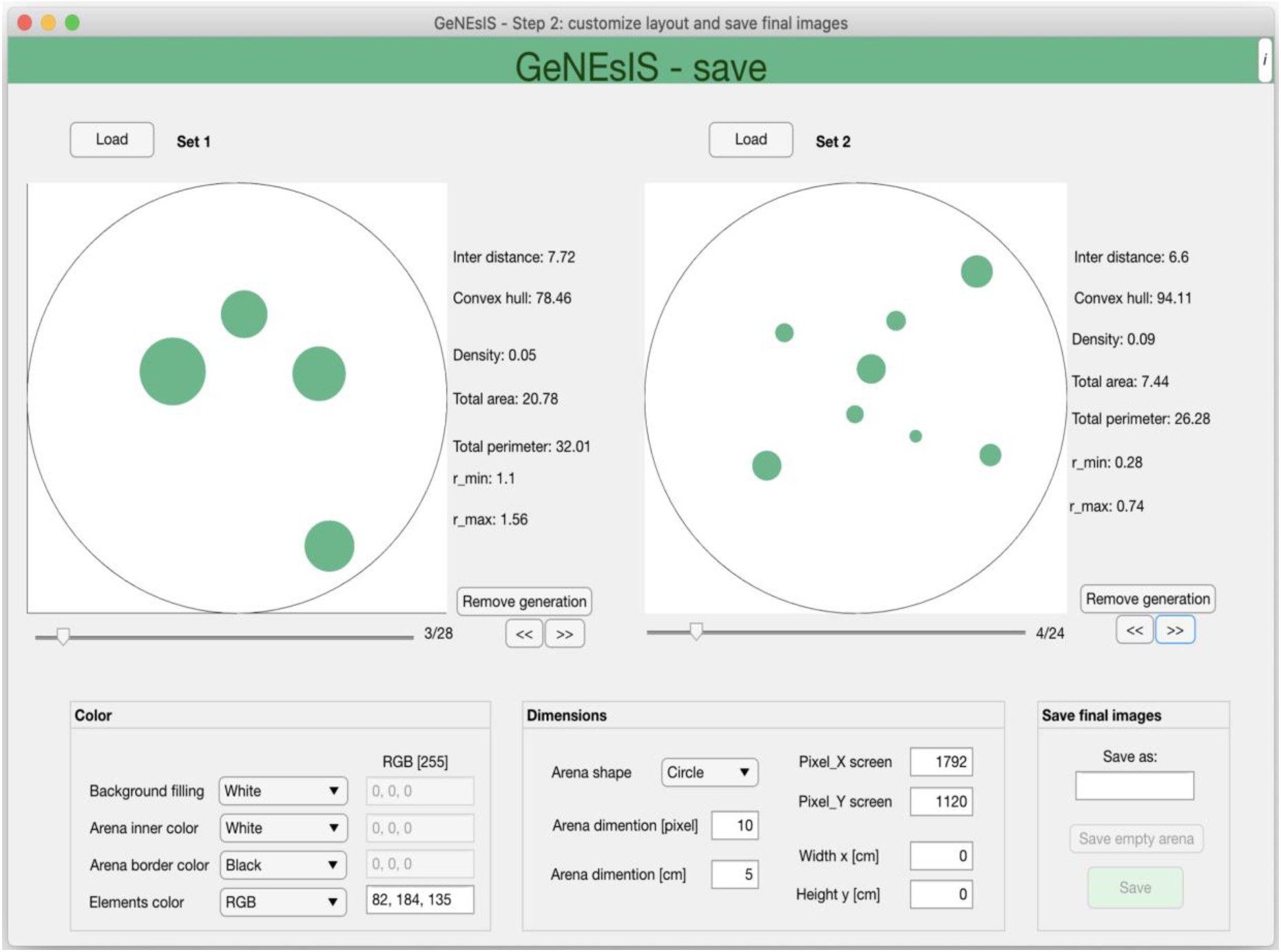
Interface for the graphic customisation of GeNEsIS stimuli, (second step, GeNEsIS_save). In this step the user can choose all the possible colours and dimensions to save the final images. The stimuli can be saved as PNG images or Matlab matrices ready to be used in a computerized presentation. The program has an intuitive user-friendly interface to guide through all the process of stimuli customization.

**Figure 4.**
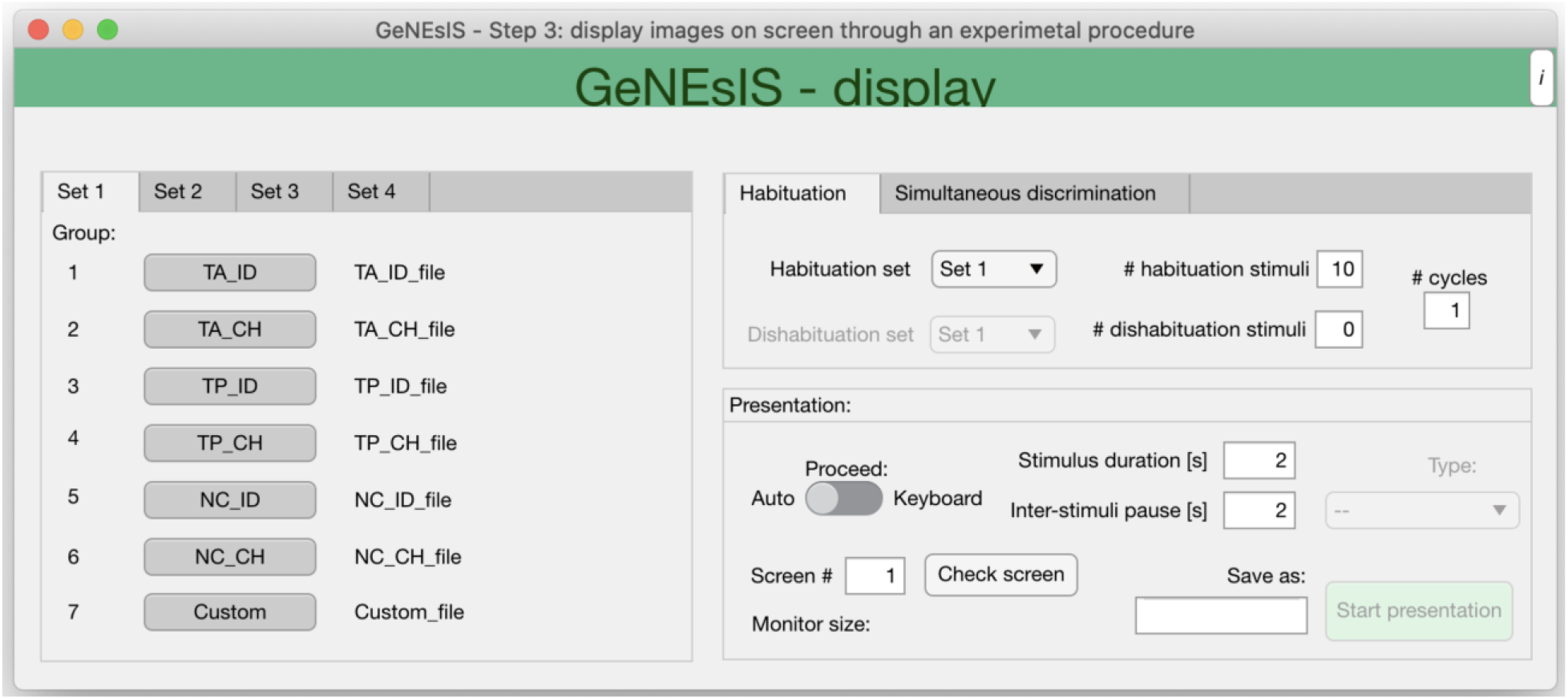
Interface for the performing of an experiment with GeNEsIS stimuli, (third step, GeNEsIS_display). This is a tool allowing two classical experimental protocols: *habituation/dishabituation* and *dual choice* tasks. The program has an intuitive user-friendly interface to guide through the whole experimental process and help in setting the presentation characteristics.

### Third step: the presentation of stimuli on screen

This step is performed with *GeNEsIS_display*. This supplementary tool, exploiting *Psychtoolbox*, allows the user to perform classical experiments on numerosity with the GeNEsIS stimuli (Figure 5).

Two type of experimental paradigms can be implemented: the “habituation/dishabituation” task and the “simultaneous dual-choice discrimination” task, which are widely used in developmental and comparative cognition (Giurfa, 2019; Halberda & Feigenson, 2008; Messina et al., 2020b; Potrich et al., 2015; Xu & Spelke, 2000). The former implies the presentation of sets of stimuli that change in all the physical properties from trial to trial, but maintaining the numerosity unchanged throughout the trials. During the dishabituation phase, novel stimuli with different characteristics (i.e., novel number of elements) are presented. The latter paradigm allows to display and visually compare two sets of stimuli that can differ in number. The stimuli to be compared can be balanced for one or more physical aspects and randomized among the session’s trials. The user can load different files created with GeNEsIS, up to four sets (numerosities) and choose the ones to present; the number of images per set is settable as well. In both experiments, the user can proceed to a subsequent trial, by pressing the keyboard or automatically, choosing the times of images presentation and pause. Again, all the sequences of stimuli presented during the experiments can be saved as an excel file containing all relevant variables during the different presentation trials, to help in the final evaluation of the experiment.

These are only two examples of experimental paradigms implemented using our program. The stimuli creation can be adapted for various needs and a large number of different experimental paradigms. Moreover, as reported above, the stimuli can be saved as image files (PNG) as well, and printed or used with other software (even different from GeNEsIS) to perform the final experiments.

## Results

### 1. Comparison with existing tools

As previously reported, many available programs have different useful characteristics, but still not all the ideal tools grouped together, for a full control over the main continuous variables. With our software we comprehend all the main useful characteristics, adding also additional features to refine the experimental design. We reported in Table 1 a sketched comparison of the main available programs developed so far (De Marco & Cutini, 2020; Gebuis & Reynvoet, 2012; Guillaume et al., 2020; Salti et al., 2017), focusing on the main tools a program could embed.

We also performed a comparison simulation with the two most recent (and most flexible, given the wider range of applications) programs (De Marco & Cutini, 2020; Guillaume et al., 2020). The outcome of the comparison is reported in Table 1. The goal of the simulation was to create different arrays for three numerosities (5, 10, 20 elements), keeping constant the convex hull (controlled at 150000 pixels) and the total area (controlled at 20000 pixels) across quantities. In order to compare the simulations at the same level and since NASCO (Guillaume et al., 2020) can create only dots with a constant radius inside an array, the array’s elements radius was kept constant in all simulations. For each numerosity we created 100 images in order to collect the final output average and relative standard error. To test the fine precision of the different software, the error tolerance was set at 0.0001% for *GeNEsIS*, or the minimum possible reached by others.

The results are reported in Figure 5, with the relative graphs representing the controlled features (convex hull and total area). In all the simulations, GeNEsIS showed a high accuracy among the numerosities created, both when the convex hull and the overall area were controlled.

It is interesting to note, across all these simulations, the very high accuracy of GeNEsIS (though the magnitudes of errors is probably negligible in most of the cases, see Figure 7).

**Figure 7.**
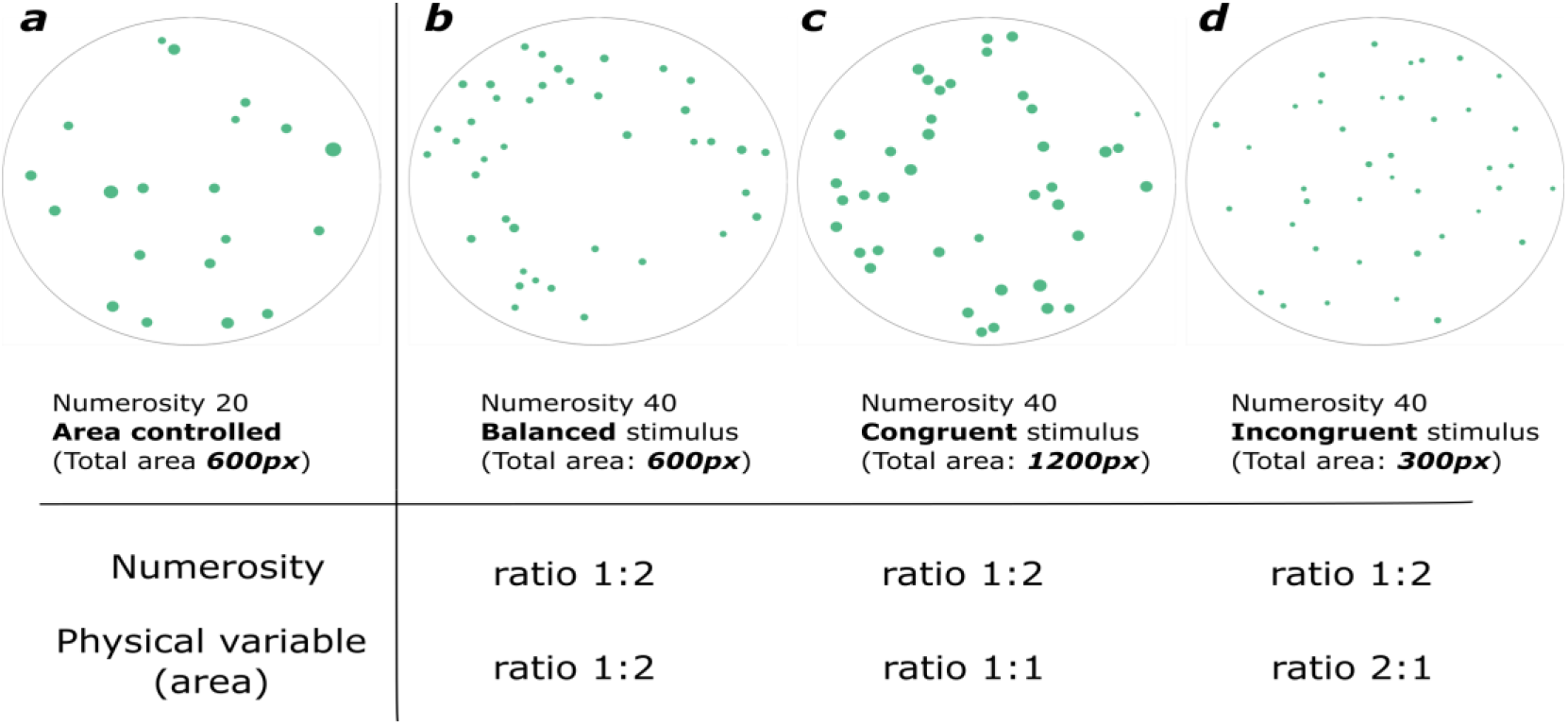
Example of comparison between 20 and 40 dots, controlled for their total area. a) Numerosity 20 with fix Total Area at 600px. b) Balanced stimulus of numerosity 40: total area is constant at 600px. c) Congruent stimulus of numerosity 40: total area (1600px) doubles as numerosity doubles. d) Incongruent stimulus of numerosity 40: total area (300px) halves as numerosity doubles.

We especially stress the usefulness of GeNEsIS for the wide range of tools it could provide compared with the other existing software (see Table 1) and the flexibility with which it can be used in many different contexts.

### 2. Examples of stimuli creation for different theoretical frameworks

#### Physical variables controlled in combination

As previously reported, GeNEsIS controls different variables independently (see Figure 1). Given the flexibility of the program the user could also control custom combinations of these variables, with the only limitations given by the geometry and visual properties of the elements’ arrays (for example it will never be possible to fix at the same time total area and total perimeter given their geometrical characterization: as the first varies with the second power of the radius, the second varies linearly). A comprehensive overview is summarized in Table 2, in which the most significant variables are considered and coupled in accordance with their geometrical limitations. An example of multiple controls between numerosities 3 and 5 is also reported (Figure 6).

Tests were additionally performed on the multiple control of different variables across numerosities, and all the main possible configurations were created. An exemplification of simulation for some of the most relevant combinations is reported in the Appendix I (Supplementary figure 1), where the accuracy of GeNEsIS in handling fixed variables and how the free variables vary across numerosities can be seen.

#### Creation of congruent and incongruent stimuli

Another possible use of GeNEsIS is given by the creation of sets of stimuli whose numerosity may vary congruently or incongruently with non-numerical physical variables. In order to report an example of this approach, we performed a simulation with 20 *vs*. 40 dots considering their total area as the continuous variable of interest. In Figure 7a the reference numerosity 20 is depicted with a fixed total area of 600 px; in Figure 7b it is reported a balanced stimulus with 40 dots, in which the numerosity doubles but the total area is still constant at 600px (overall surface are is equal between the two numerosities). If instead one would create a congruent stimulus, this can be done increasing the total area as the numerosity increases (Figure 7c: the total area doubles with numerosity); on the contrary, it is possible to create incongruent stimuli by decreasing the overall area as the numerosity increases (Figure 7d shows a stimulus incongruent for the total area).

This idea can be applied to every kind of variable and adapted to different theoretical backgrounds, showing the huge flexibility of GeNEsIS.

### Discussion and conclusions

Numerical abilities are important for many different species, since organisms use numerical information to perform adaptive choices (Haun et al., 2010; Nieder, 2020). However, researchers also identify alternative strategies to number encoding, based on perceptual cues that vary according with numerosity (Gebuis & Reynvoet, 2011, 2012); this raised the hypothesis of the existence of a more general *sense of magnitude* (Leibovich et al., 2017). Indeed, organisms could use the variation in continuous cues, such as area, perimeter or space occupied by elements, to discriminate arrays containing a different number of items.

For these reasons, a careful consideration of the variation in continuous physical variables when visual arrays are presented should be a mandatory aspect of investigation in numerical cognition. Despite previous studies having carefully applied several controls in the investigation of numerical abilities of different species, a common and standardized experimental method to guide such controls seems to be missing.

In the last years, new programs were developed to help researchers with numerosity elements array creation (De Marco & Cutini, 2020; Guillaume et al., 2020; Salti et al., 2017). Despite their usefulness, each of them lacks some fundamental tool to allow full flexibility in creating controlled stimuli (see Table 1 for a comparison between programs). Here we propose GeNEsIS as a tool to create numerousness stimuli controlled in their physical continuous variables in a standardized and highly accurate (Figure 5 and Appendix I: Supplementary Figure 1) way. Our program sums up all the aforementioned utilities in a comprehensive tool, implementing the until now missing features (Table 1). In particular with GeNEsIS the user can easily control convex hull, mean inter distance, density, total area, total perimeter, elements’ number and size, independently or in combination, with the only limitations imposed by the geometry (Table 2). Moreover, the layout is completely customizable, allowing also to create elements with different colours and shapes (circles, triangles, diamonds and squares). All the procedures are guided by an intuitive graphical user interface, permitting even the people less practical with coding to easily create the stimuli. In addition, we also implemented a tool to perform the presentation of these images on screen, during standardized experiments of habituation/dishabituation or dual choice tasks.

With this program we hope to fill the gap in the current state of the art in numerical cognition experiments, providing a tool that can improve the way in which experiments are conducted, reducing at a minimum the bias in numerousness array due to continuous physical variables, and standardizing the procedure of stimuli creation and presentation to facilitate experimental replicability and inter and intra-species comparisons.

## Additional information

## Availability of materials and code

All the reported materials, data and the program code are freely available at https://github.com/MirkoZanon/GeNEsIS.

## Funding

This work was supported by a European Research Council Grant to G.V. (ERC grant 833504-SPANUMBRA) and by Progetti di Rilevante Interesse Nazionale (PRIN 2017 ERC-SH4–A 2017PSRHPZ)

## Competing interests

The authors declare no competing interests.

## Authors’ contributions

M.Z. designed and implemented the program. M.Z., D.P., M.B. contributed in testing the program. D.P, M.B., M.Z contributed to the writing of the manuscript. G.V. supervised the project. All the authors contributed to the final version of the manuscript.

## Appendix I Supplementary

**Supplementary figure 1:**
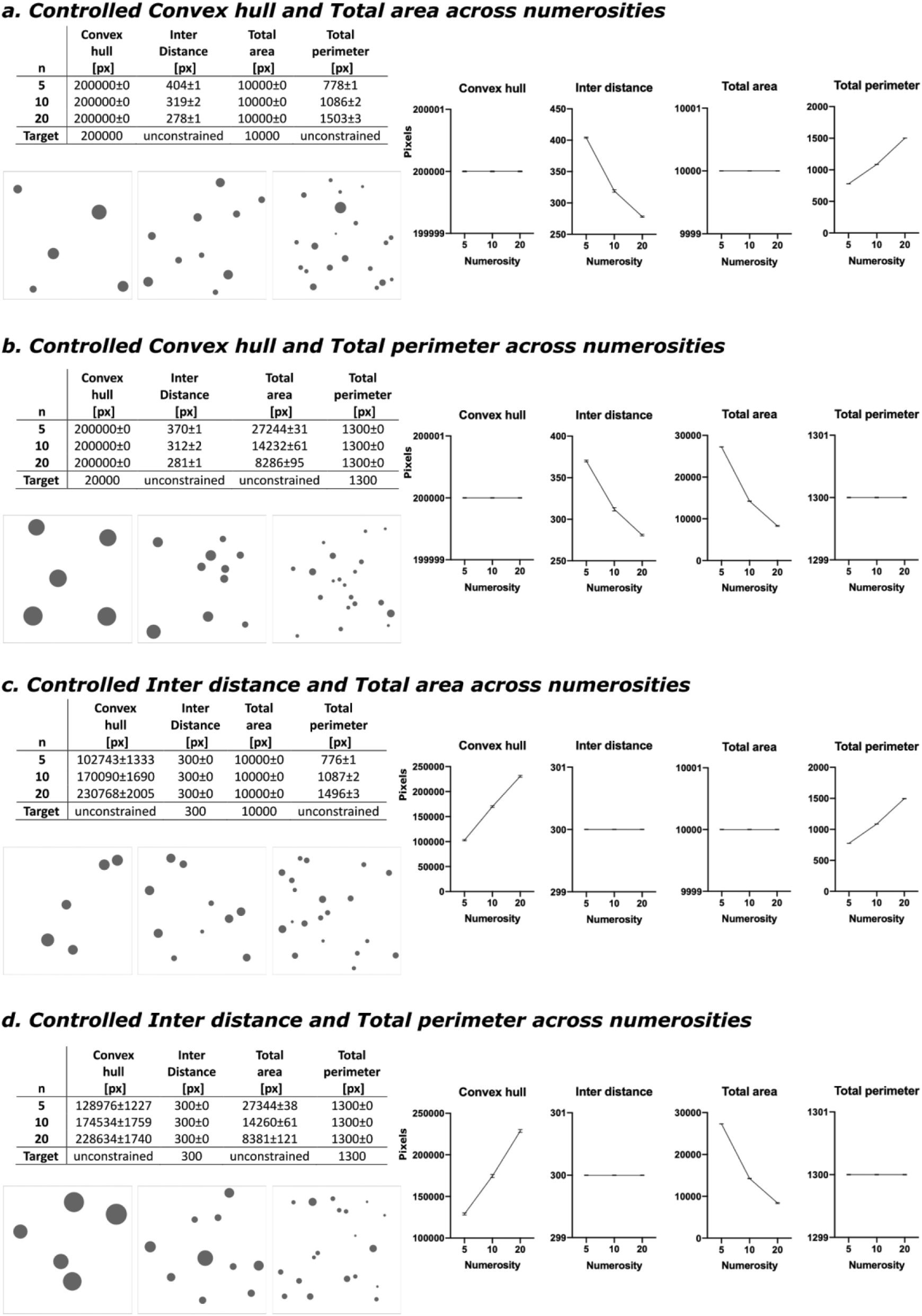
Example of the main combinations of variables controlled at a time, across different numerosities (5,10 and 20 elements). Four different combinations were tested (a. CH 200000px + TA 10000px; b. CH 200000px + TP 1300px; c. ID 300px + TA 10000px; d. ID 300px + TP 1300px). For each combination and each numerosity 100 images were created: results show the mean and standard error over these groups. In order to fine test the precision of GeNEsIS a small tolerance error was set (0.001%). The output mean results for the 4 main variables considered here are reported in the tables and in the relative graphs. One image per numerosity is also shown.

## Notes

https://github.com/MirkoZanon/GeNEsIS

### Competing Interest Statement

The authors have declared no competing interest.

